# Mathematical modelling of molecular heterogeneity identifies novel markers and subpopulations in complex tumors

**DOI:** 10.1101/283903

**Authors:** Lulu Chen, Niya Wang, Robert Clarke, Zhen Zhang, Yue Wang

## Abstract

Intratumor heterogeneity, as both a major confounding factor and an underexploited information source, is widely implicated as a key driver of drug resistance. While a handful of reports have demonstrated the potential of supervised methods to deconvolute intratumor heterogeneity, these approaches require a priori information on the marker genes or composition of known subpopulations. To address the critical problem of the absence of validated marker genes for many (including novel) subpopulations, we developed convex analysis of mixtures (CAM), a fully unsupervised deconvolution method, for identifying marker genes and subpopulations directly from original mixed molecular expressions.

## I. INTRODUCTION

Drug resistance is a major barrier to increasing overall survival rates, whether this resistance is present at the time of presentation or emerges after treatment response. Multiscale intratumor heterogeneity (ITH) is characteristic of complex tumors and is widely implicated as a key driver of drug resistance. Precisely how ITH develops within and across scales is unclear.

ITH, arising from multiple subpopulations within a tumor, is both a major confounding factor in studying individual subpopulations and an underexploited information source for characterizing complex tumors [1]. ITH is multiscale, i.e., evident at many levels (e.g., genetic, transcriptomic, proteomic, metabolic, cellular, phenotypic). Complex tumors can be characterized by the identity, composition, and expression profile of possibly unknown subpopulations [2], where subpopulations are often defined by marker genes (genes whose expressions are exclusively enriched in a particular subpopulation [3, 4]). While many reports have demonstrated the potential of supervised methods to deconvolute ITH, these approaches almost exclusively require a priori information on the known subpopulations [2, 3]. Such supervised methods (including cell sorting) have difficulty dissecting ITH that are subtle, condition-specific or previously unknown [1, 5].

To address the critical problem of the absence of validated marker genes for many (including novel) subpopulations, we recently developed a fully unsupervised computational method (convex analysis of mixtures – CAM) [6, 7] that can identify subpopulation-specific marker genes directly from the original mixed expressions - a nontrivial task. CAM requires no prior information on the number, identity, or composition of the subpopulations present in mixed samples [8], and does not require the presence of pure subpopulations in sample space [9, 10]. CAM dissects heterogeneity into molecularly distinct subpopulations by leveraging the advantages of both tissue-wide and single-cell approaches [10, 11]. Discerning differences among single cells can provide valuable information about inter-cellular heterogeneity but may lose critical information of cell-cell interactions and is prone to cell-cycle/state confounders; while sample-wide measures give a detailed picture of the average state of a cell population but at the cost of losing information about inter-subpopulation heterogeneity.

## II. RESULTS

### A. Validation of CAM on real dataset

CAM’s ability for unsupervised data deconvolution has been validated on several real benchmark gene expression datasets (GSE19830, GSE11058) [6]. More recently, CAM analysis of proteomic data from left anterior descending coronary artery (LAD) and abdominal aorta (AA) specimens from a large cohort identified 58 informative proteins of normal (NL), fatty streak (FS), and fibrous plaque (FP) burden levels with 48 proteins verified in an independent validation cohort [12].

### B. Application of CAM to model ITH in breast cancer

To addressed ITH remodeling in the context of endocrine resistance, the MCF-7 variants LCC1 cells (estrogen independent; antiestrogen sensitive) and LCC9 cells (estrogen independent; antiestrogen resistant) are mixed in different ratios, and treated with drug ICI or vehicle. Hierarchical clustering of their iTRAC proteomic data shows that the molecular profiles of 1:1 and 5:1 (LCC1:LCC9) ICI-treated and 1:1 untreated mixtures cluster with LCC9 (Fig. 3a). Untreated 5:1 mixtures are close to untreated LCC1. Data from the mixtures were deconvoluted in silico (unsupervised) by CAM into two subclone signals (C1, C2; Fig. 3b). When combined with data from pure cells, PCA shows that C1 signals from untreated 1:1 and 5:1 are close to LCC1, and the C2 signals from ICI treated and untreated mixtures are close to LCC9. C1 signals from the treated 1:1 and 5:1 mixtures form a separate cluster distant from LCC1 and LCC9 cells, implying that resistant cells and sensitive cells alter each other’s molecular signatures. These reflect the importance studies into the in vivo tumor microenvironment.

## III. METHODS

### A. Mathematical foundation

Consider a set of *J* heterogeneous samples of varying composition of unknown *K*(≤ *J*) subpopulations. We assume that the measured expression level is linearly proportional to the abundance of that subpopulation (without log transformation [13], Fig. 1). We further assume that gene expression values are non-negative and adopt the definition of subpopulation-specific marker genes as those genes whose expression values are exclusively enriched in a particular subpopulation [3] (Fig. 1a). Fundamental to the success of CAM is a novel insight that subpopulation-specific marker genes that define latent pure subpopulations reside at the extremities of the scatter simplex formed by all genes, while the interior of the simplex is occupied by co-expressed genes (whose values are linear non-negative combinations of pure subpopulation expression values) (Fig. 1b,1c). We can then identify novel marker genes by geometrically locating the vertices of the multifaceted simplex that most tightly encloses the gene expression profiles and has the same number of subpopulations as vertices.

**Figure 1.**
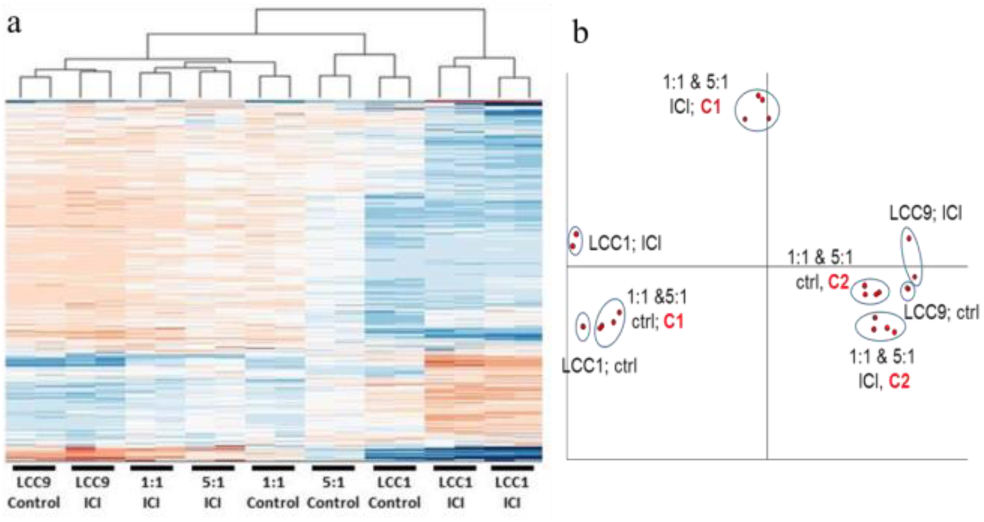
Clustering (a) and Principal Component Analysis (b) plots of LCC1/LCC9 mixed cells using iTRAC data (proteome); Ratios=LCC1:LCC9; ICI=100 nM; ctrl=vehicle.

**Figure 2.**
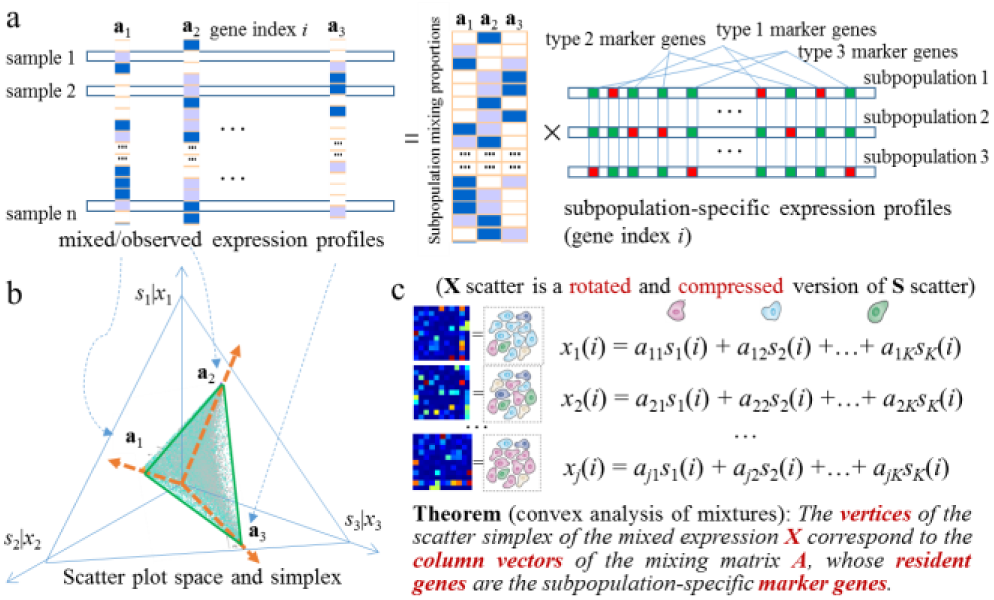
CAM principles for unsupervised identification of novel subpopulation-specific marker genes

### B. Data preprocessing

First, we eliminate genes whose signal intensity (vector norm) is lower than 5% (noise) or higher than 95% (outlier) of the mean value over all genes. The signals from these genes are unreliable and could have a negative impact on the subsequent analyses. Second, when J >> K, dimension reduction is performed on the raw measurements using principal component analysis, sampling clustering or nonnegative matrix factorization techniques, to improve the efficiency of subsequent analyses [5]. Third, a sum-based standardization is applied to gene vectors (expression values across samples) to generate scatter simplex. Forth, we aggregate gene vectors into representative clusters using affinity propagation clustering (APC) [14] to further reduce the impact of noise/outlier data points (gene expression vectors over samples) and improve the efficiency of CAM.

### C. Convex analysis of mixtures (CAM)

To identify the vertices of clustered convex set *X* (scatter simplex of mixed expression profiles), we performed CAM on the obtained *M* cluster centers {*g*_*m*_} of gene vectors. We assumed *K* true vertices and conducted an exhaustive combinatorial search (with total 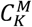 combinations), based on a convex-hull-to-data fitting criterion, to identify the most probable *K* vertices. We used the margin-of-error

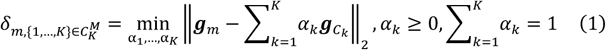

to quantify the distance between *g*_*m*_ and convex set *X* defined by {*g*_*k*=1,…,*K*_}, where we have zero margin-of-error if {*g*_*m*_} is inside *X*. We then selected the most probable K vertices when the corresponding sum of the margin-of-error between the convex hull and the remaining “exterior” cluster centers reaches its minimum: Subsequently, we identified the indices of subpopulation-specific marker genes based on the memberships associated with *K* vertices.

### D. Model selection

We used MDL, a widely-adopted and consistent information theoretic criterion [15], to automatically detect the number K of subpopulations in the heterogeneous samples by minimizing the total description code length:

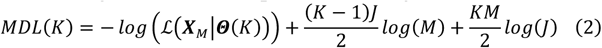

where ℒ (∙) denotes the joint likelihood function of the clustered latent variable model, *X*_*M*_ denotes the set of *M* gene vector cl0075ster centers, and Θ(*K*) denotes the set of freely adjustable parameters in the clustered latent variable model.

### D. Software

A Java-R package of CAM is available at http://mloss.org/software/view/437, providing comprehensive analytic functions and graphic user interface (GUI) to help users readily apply CAM method to their own datasets. Data preprocessing (noise/outlier removal and dimension reduction) needs to be performed by users manually.

### E. Areas where this method is useful

As a non-negative blind source separation method, CAM can be applied to deconvolute underlying sources of various data types, not only mRNA and proteomics data mentioned in this paper but also microRNA, methylation, magnetic resonance imaging data, etc.

